# Deep learning of gene relationships from single cell time-course expression data

**DOI:** 10.1101/2020.09.21.306332

**Authors:** Ye Yuan, Ziv Bar-Joseph

## Abstract

**Motivation:** Time-course gene expression data has been widely used to infer regulatory and signaling relationships between genes. Most of the widely used methods for such analysis were developed for bulk expression data. Single cell RNA-Seq (scRNA-Seq) data offers several advantages including the large number of expression profiles available and the ability to focus on individual cells rather than averages. However, this data also raises new computational challenges.

**Results:** Using a novel encoding for scRNA-Seq expression data we develop deep learning methods for interaction prediction from time-course data. Our methods use a supervised framework which represents the data as a 3D tensor and train convolutional and recurrent neural networks (CNN and RNN) for predicting interactions. We tested our Time-course Deep Learning (TDL) models on five different time series scRNA-Seq datasets. As we show, TDL can accurately identify causal and regulatory gene-gene interactions and can also be used to assign new function to genes. TDL improves on prior methods for the above tasks and can be generally applied to new time series scRNA-Seq data.

**Availability and Implementation:** Freely available at https://github.com/xiaoyeye/TDL.

**Supplementary information:** Supplementary data are available at *XXX* online.

## 1 Introduction

Inferring gene relationships and function from expression data has been a major focus of computational biology over the last two decades (Marbach, et al., 2012; Stuart, et al., 2003). While many of these methods were developed for, and applied to, static data, time series data may be even more appropriate for such analysis. By profiling genes over time researchers can identify not just correlations (that may imply interactions) but also causation (Finkle, et al., 2018). Indeed, several methods have been developed and used to infer causal interactions from time series gene expression data. Some of these methods were originally developed for static data and later applied to time series as well (Huynh-Thu and Geurts, 2018; Huynh-Thu, et al., 2010). Other methods were specifically developed for time series data (Kim, et al., 2011). These methods ranged from alignment methods (Qian, et al., 2001) to regression analysis (Huynh-Thu and Geurts, 2018), methods based on granger causality inference (Finkle, et al., 2018; Kim, et al., 2011) and methods utilizing various forms of graphical models (Schulz, et al., 2012; Zou and Conzen, 2005).

All of the methods discussed above were developed for bulk expression analysis. Recently, researchers have been performing time series studies using single cell RNA-Sequencing (scRNA-Seq) (Mayer, et al., 2018; Semrau, et al., 2017; Shin, et al., 2019; Soldatov, et al., 2019). While such studies provide much more detailed information about the genes and cell types involved in the process, they also raise new challenges. A major issue for using such data to infer gene relationships is the fact that cells profiled in a specific time point cannot be tracked. Thus, it is not clear which cell in the next time point is a descendent (or closely related) to a specific cell in the previous time point making it hard to determine exact trajectories for genes. Another challenge arises from the large number of cells being profiled at each time point. Finally, the fact that cells in a time point may be from several different types and may not be fully synchronized (Wang, et al., 2017) makes it harder to establish a specific pattern for temporal analysis.

To address these issues some methods first perform pseudo-time ordering of the cells (Trapnell, et al., 2014) followed by regression or correlation analysis (Specht and Li, 2017). While such methods can indeed identify some of the casual interactions, they depend very strongly on specific assumptions including the accuracy of the ordering achieved, the expected time lag (in regression) or the format of the interaction (in the case of correlation).

We have recently developed the convolutional neural network for coexpression (CNNC) analysis method which is focused on inferring gene-gene relationships from static scRNA-Seq data (Yuan and Bar-Joseph, 2019). Unlike prior methods for inferring interactions, CNNC does not make any assumptions about the specific attributes or correlations that can be observed for interacting genes. Instead, it uses a supervised framework to train a CNN for predicting such interactions. The main novel idea of CNNC is the way gene expression data is represented. While most methods represent such data as vectors (for a single cell) or matrix (for a collection of cells) CNNC converts expression data into an image like histogram representation. This enables CNNC to take full advantage of the ability of CNNs to utilize local substructures in the data.

While CNNC can be directly applied to time series scRNA-Seq data, such application ignores the temporal information and so does not lead to optimal results as we show. Here we present two extensions of CNNC and show that by explicitly considering the temporal information we can greatly improve the performance of the method. The first extension we consider is converting the input from a 2D to a 3D representation of the joint probability function over time, while still using a CNN. The second uses a different type of deep NN termed Long Short Term Memory (LSTM) which is specifically appropriate for time series analysis (Xingjian, et al., 2015). By using LSTM we can encode both, an image for each time point and the way the images are related over time enabling the method to infer interactions both within and across time. We term the new methods we develop Time-course Deep Learning (TDL). We discuss how to formulate TDL models, how to train them and learn parameters for them and how to apply them to time series scRNA-Seq data. Testing the methods on several recent time series scRNA-Seq datasets we show that they can accurately infer interactions, causality and assign function to genes and that TDL improves upon current and prior methods suggested for these tasks.

## 2 Methods

To utilize the advantages of convolutional neural networks (CNNs) we encode time series scRNA-Seq data using a 2D or 3D histogram termed normalized empirical probability density function (NEPDF). These inputs encode, for each pair of genes, the co-occurrence frequency of their values either across cells in all time points (2D NEPDF) or for all cells in a specific time point (3D NEPDF). Next, we use these as features and train a Time-course Deep Learning (TDL) model using a supervised computational framework to predict causality, infer interactions and assign function to genes.

### 2.1 Data used

To test the ability of our TDL methods to predict interaction and causal relationships we used time series scRNA-Seq data from four mouse and human embryonic stem cell studies (mESC and hESC). Data was downloaded from the accession numbers of GSE79578, GSE65525, E-MTAB-3929 and GSE75748 (Chu, et al., 2016; Klein, et al., 2015; Petropoulos, et al., 2016; Semrau, et al., 2017). The two mESC datasets profiled 3,456 and 2,718 cells with nine and four time points, respectfully. The hESC datasets profiled 1,530 and 758 cells, both in six time points (we removed the first time point for the 1^st^ hESC dataset since it contains much less cells than other time pints). We also used a mouse brain time-course single cell data (cortex cells with 3 time points (GSE104158) (Mayer, et al., 2018)) when testing our method’s ability to infer new functional genes. Count expression data was normalized so that all cells had the same total value of 10,000. Some of the datasets provided RPKM data, so we adopted it directly. Genes that are never expressed were filtered out.

### 2.2 Ground truth interactions

TDL rely on a supervised learning framework so need both positive and negative inputs for training. We used TF-gene interaction information to construct the skeleton of gene regulation networks and then defined the edges as positive pairs for training and testing for both causality and interaction prediction tasks. To determine TF’s target genes we used mESC and hESC ChIP-seq data. We relied on a strict peak p-value cutoff to identify binding sites (p value < 10^-400^ based on MACS2 (Zhang, et al., 2008)). We next defined a ‘promoter region’ as 10Kb upstream and 1Kb downstream from the TSS of each gene and assigned TFs to regulate genes if a peak for that TF was identified in the promotor region for the gene as has been previously done (Schulz, et al., 2013; Yuan and Bar-Joseph, 2019). Genes whose promoter region had a significant peak were regarded as positive targets, while those that did not were used as the negative set. To test the accuracy of the functional gene assignments we downloaded from GSEA (Subramanian, et al., 2005) 680 cell cycle, 93 rhythm, 184 immune and 137 proliferation genes.

### 2.3 2D and 3D NEPDF input construction

The input to our TDL methods is a 3D image-like NEPDF. For a gene pair (*a, b*), 2D and 3D NEPDF are generated as follows: Expression ranges for all genes across all cells are divided into 16 equal bins in log space. Next, expression values of gene pairs in each cell were used to compute a 16×16 matrix where each entry (*i, j*) represents the co-occurrence frequency of gene pair (*a, b*) with the *i*th, and the *j*th expression level, respectively. Due to dropout in scRNA-seq data, the value in the zero-zero position always dominates the entire matrix. To solve this problem, one more log transformation with a psedudo-count was applied to every entry to generate the final 2D normalized matrix. A 3D NEPDF is generated in a similar way by constructing a separate 2D NEPDF for each of the time points in the input data. For the 3D NEPDF we used an 8×8 matrix for each time point rather than 16×16 since the number of cells in each time point is much smaller than the total number of cells. Thus, the 3D NEPDF is a tensor with dimension of #time point (T)×8×8. We note that while the dimensions listed above are the ones used in the paper, we tried other dimensions as well for both the 2D and 3D representations. To select the optimal dimension for each method we performed analysis of various potential input sizes, and determined that the optimal size for TDL model is #time point (T)×8×8 and that the optimal input size for CNNC is 16×16. See **Fig. S6** and **S7** for details. In the 3D tensor, each column (row) represents gene *a*(*b*)’s expression level, while each depth represents the time point *t* in biological experiment settings. As a result, the entry in the 3D NEPDF represents the normalized frequency (probability) of co-occurrence of its corresponding gene pair expression in time point *t*. **Fig. 1**, presents an overview of the encoding. See Supporting Methods for details about constructing 2D and 3D NEPDF, normalization of the histogram data and transformations used to overcome dropouts and the large concentration that is often observed at the [0,0] point.

**Fig. 1.**
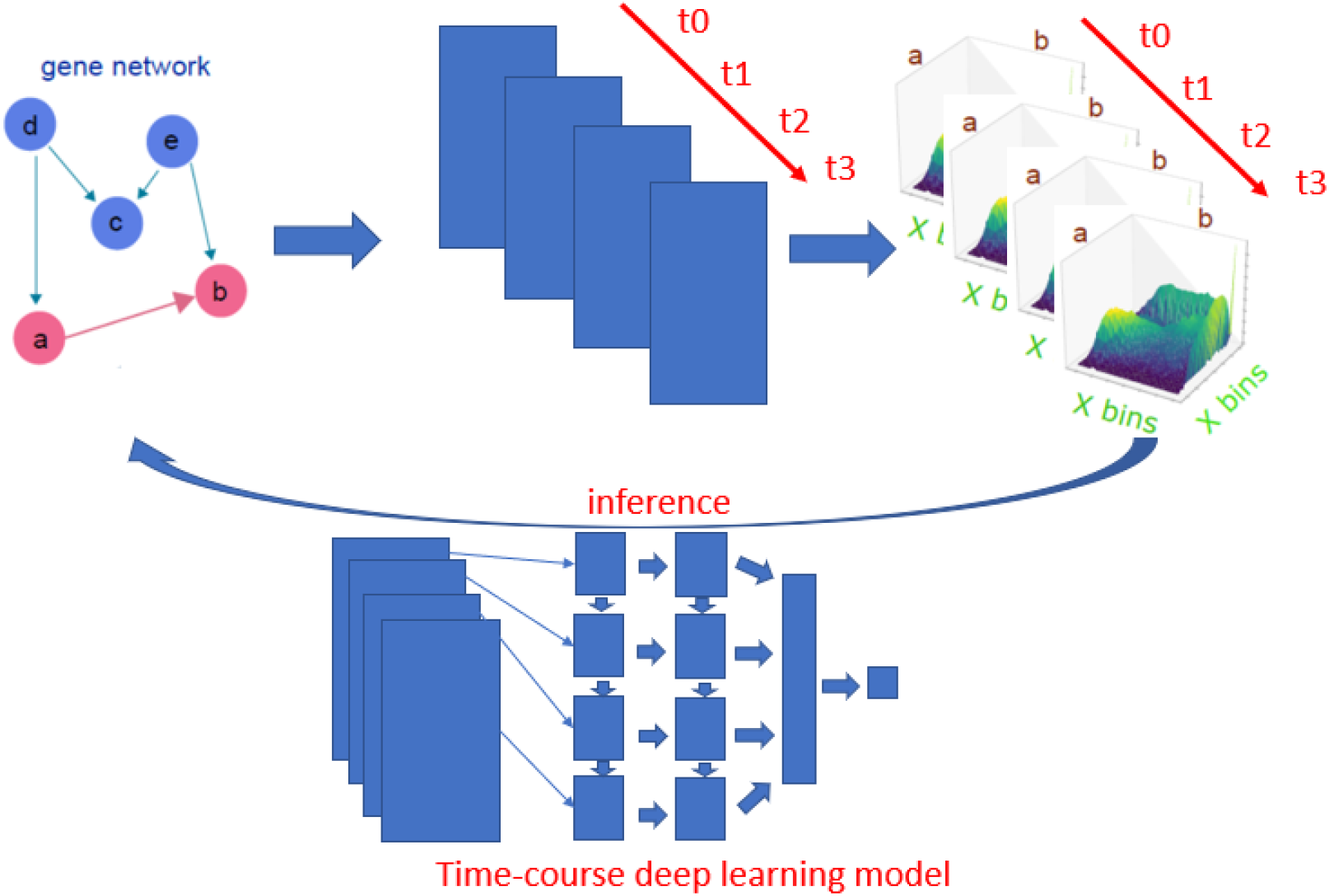
Time-course deep learning (TDL) model architecture. To infer gene interactions (top left) we first convert time-course single cell expression data to a 3D tensor, which we term normalized empirical probability distribution function (NEPDF). Each 2D slice of the NEPDF captures the co-expression of a pair of genes at one of the time points profiled and the 3D NEPDF represents their co-expression over time. 3D NEPDF is then used as input to atime-course deep learning (TDL) model. The model is trained using labeled positive and negative pairs. The figure shows the convolutional Long short-term memory (LSTM) architecture which is one of the two TDL models we tested. This mode consists of two stacked LSTM layers, followed by a dense layer which concatenates all convolutional hidden state from LSTM layer and then a final output (classification) layer. See **Fig. S1** for the other TDL architecture we tested, 3D CNN.

### 2.4 Architectures for TDL models

While our previous models can use the 2D representation (Yuan and Bar-Joseph, 2019) this representation does not utilize the temporal information. We have thus developed two TDL models that work directly on the 3D tensor input. The output of all models is a *N*-dimension (*N*D) vector, where *N* depends on specific task on which the TDL is trained (for example, *N*=1 for interaction predictions and 3 for interaction and causality predictions). The first TDL model we consider is termed 3D CNN and is a direct extension of the 2D CNNC method (Yuan and Bar-Joseph, 2019). In general, 3D CNN consists of one T×8×8 tensor input layer, several intermediate 3D convolutional layers, Max pooling layers, one flatten layer and a *N*D classification layer (see **Fig. S1** for structure details). While CNNs can utilize time series data if the data is encoded as a 3D tensor, they are not intended for time series or sequential data. Another type of neural network, termed Recurrent neural network (RNN) (Mikolov, et al., 2010; Min, et al., 2017) is more appropriate for such data model. In RNNs, the activation function used for subsequent inputs (for example, later time points) uses the values learned from previous inputs (earlier time points) (**Fig. 1**). Specifically, for time *t* the network computes the following value for the hidden layer:

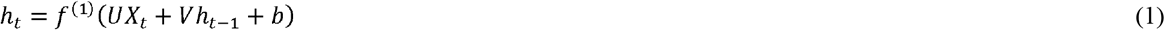

where *h*_*t*−1_, is a hidden state computed for the previous time point and *X_t_* is the input for time t. *U* and *V* are (learned) weight matrices, *f*^(1)^ is an activation function and *b* is bias term. We compute the output by setting:

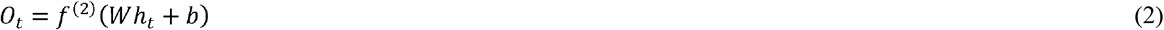

Where *O_t_* is output at time t, and *f*^(2)^ is an activation function. See **Fig. S2** for details on the RNN architecture and its unrolled sequential structure.

While successful, RNN can suffer from gradient vanishing or gradient explosion problems (Hochreiter, 1998). A popular way to overcome these problems is to use a variant of RNN termed Long Short-Term Memory (LSTM (Hochreiter and Schmidhuber, 1997)). LSTMs utilizes a cell memory unit, and combines the current (time point) input, the previous cell memory and the computed hidden state to update the cell memory. An output gate is used to control the information propagation from cell memory to hidden state.

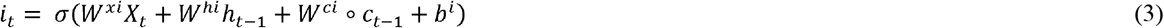

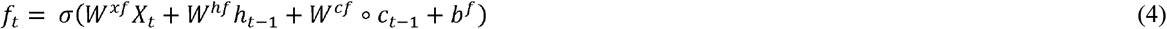

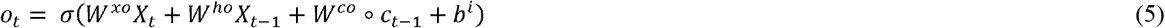

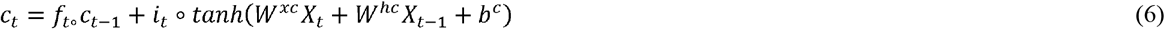

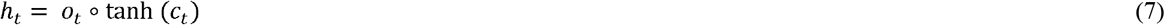

where *σ* is sigmoid function, *i_t_, f_t_, o_t_, c_t_, h_t_* are input gate, forget gate, output gate, cell and hidden state vectors at time *t*, and ° represents Hadamard product. *h_t_* serves as both, input to the next layer over time and the output for the final state in the model.

The original LSTM method uses a fully connected NN. However, for our, time-course 3D NEPDF we also need to account for spatial information encoded for each time point. As a result, we use a convolutional version of LSTM. The conv-LSTM uses the following functions:

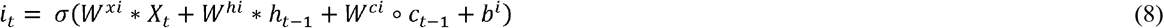

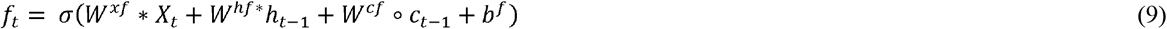

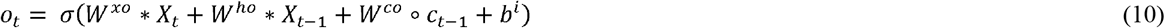

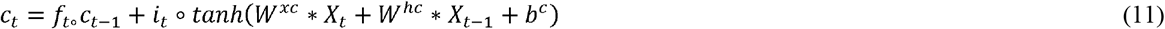

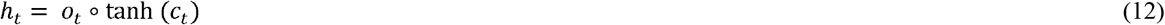

Where *σ* is sigmoid function, and *i_t_, f_t_, o_t_, c_t_, h_t_* are input gate, forget gate, output gate, cell vector at time t, and * represents convolution operator. All hidden states are then converted as a vector followed by classifier. See **Figs. S3** and **S4** for complete structure details of conv-LSTMs we used.

### 2.5 Train and test strategy

We used three-fold cross validation strategy to train and test models for all tasks. We separated the data to avoid information leakage: For the gene causality and interaction prediction tasks, the generated NEPDF was separated based on the TFs. In other word all predictions related to a specific TF were either used only in training or only in testing. In each fold we computed AUROC for each left-out TF and then the curves and values for TFs in each fold were combined in the figures presented. For gene function assignment, NEPDFs for training did not use any of the test genes to avoid information leakage. We randomly selected genes not in the known (positive) function gene set as negative genes. Positive (negative) gene were split into positive (negative) training and test gene set. For the train (test) set, all possible gene pairs where the 1^st^ gene is in positive training set and the 2^nd^ is in positive train (test) set were treated as positive pairs, and pairs where the 1^st^ gene is in the positive training set and the 2^nd^ from the negative train (test) set are treated as negative. All the deep learning models were trained with the stochastic gradient descent (SGD) algorithm using a maximum training epoch of 100. Accuracy of the internal validation set was monitored for early stopping. See **Tab. 1** for runtime information based on the size of the input and **Tab. S1** for the time of all models for interaction task.

**Tab 1.** Run times (second per epoch) for deep learning methods for the hESC1 dataset.

| Input size | 3D CNN | Deep conv-LSTM | 2D CNNC |
| --- | --- | --- | --- |
| 5×4×4 | 10 | 24 | - |
| 5×8×8 | 35 | 57 | - |
| 5×16×16 | 177 | 360 | - |
| 16×16 | - | - | 9 |

## 3 Results

### 3.1 Causality predictions using TDL models

One of the major advantages of time series when compared to static expression data is the ability to use lagged relationships between expression profiles to infer the direction (causality) of gene-gene interactions (Bar-Joseph, et al., 2012; Qian, et al., 2001). We thus first tested if our TDL models can correctly infer the direction of interaction for TF-gene pairs since for these we know the regulator and the direction. For this we used two mouse embryonic stem cell (mESC) and two human embryonic stem cell (hESC) time-course SC datasets and mESC, and hESC ChIP-seq data from the Gene Transcription Regulation Database (GTRD) (Yevshin, et al., 2017). We used 38 and 36 TFs for testing the two mESC expression datasets due to differences in coverage of the two datasets. For both hESC data, we used 36 TFs. We trained all models with the following labeled data: A TF *a* and its target gene *b*, (*a, b*) will be assigned a label of 1 while the label for (*b, a*) is 0. In other words, the goal is to use the time series data to infer not which genes are interacting but rather the direction of the interaction.

We compared the TDL approaches (3D CNN and Conv-LSTM) with a 2D CNNC (Yuan and Bar-Joseph, 2019) (which is also supervised), and unsupervised methods that utilize granger causality and lagged regression analysis to infer the interaction direction (Kim, et al., 2011). The granger causality inference method was applied to 3 of the 4 datasets since the 2^nd^ (mESC2) only had 4 time points which turned out to be less than the minimal number of time points required by the method. We tested two versions for the granger causality method. The first strategy uses the average expression of genes in each time point for the analysis, while the second first orders all cells using on the pseudo-time ordering determined by Monocle3 (Cao, et al., 2019) and based on this order performs the regression analysis. We note that we were only able to obtain useful results when using Monocle3 for two of the four datasets (mESC1 and hESC2) and so did not perform ordered analysis for the other two datasets (see **Fig. S5** for the outcome of pseudotime ordering for all four datasets). We did not compare causality prediction with other unsupervised methods (Pearson correlation and MI) since these are symmetric and so cannot infer directionality information. Three-fold cross validation was used to evaluate all methods. Since each fold contains more than 10 TFs, area under receiver operating curve (AUROC) was calculated for each TF, and the curves for each TF were then collected to compute the final performance for each fold.

Results presented in **Figs. 2A** and **S8** show that pseudo-time ordering improved the granger causality analysis results though using such ordering results for this method is still inferior to deep learning methods. We also observe that the TDL models outperform 2D CNNC on all the datasets (mean AUROC for mESC1, 0.73 vs 0.72; mESC2, 0.67 vs 0.61; hESC1, 0.77 vs 0.73; and hESC2, 0.66 vs 0.65). We also plotted the input representation for a few correctly predicted causal pairs. As expected (**Fig. 2B-E**), we observe that a shift or phase delay is always from the affecting to the affected gene. Unlike prior methods that require that they user specifies the lag duration, TDL determines the importance of such shift and the length based on training information only, highlighting the flexibility of the representation and method. We also explored sample that was only correctly assigned by TDL while the 2D CNNC incorrectly inferred the wrong direction for them (**Fig. 2F-H**). We found that the model trained with the 2D NEPDF (CNNC) focuses on low expression (marginal) regions, while TDL models focus on temporal dynamics including phase differences which enables them to make the correct assignments.

**Fig. 2.**
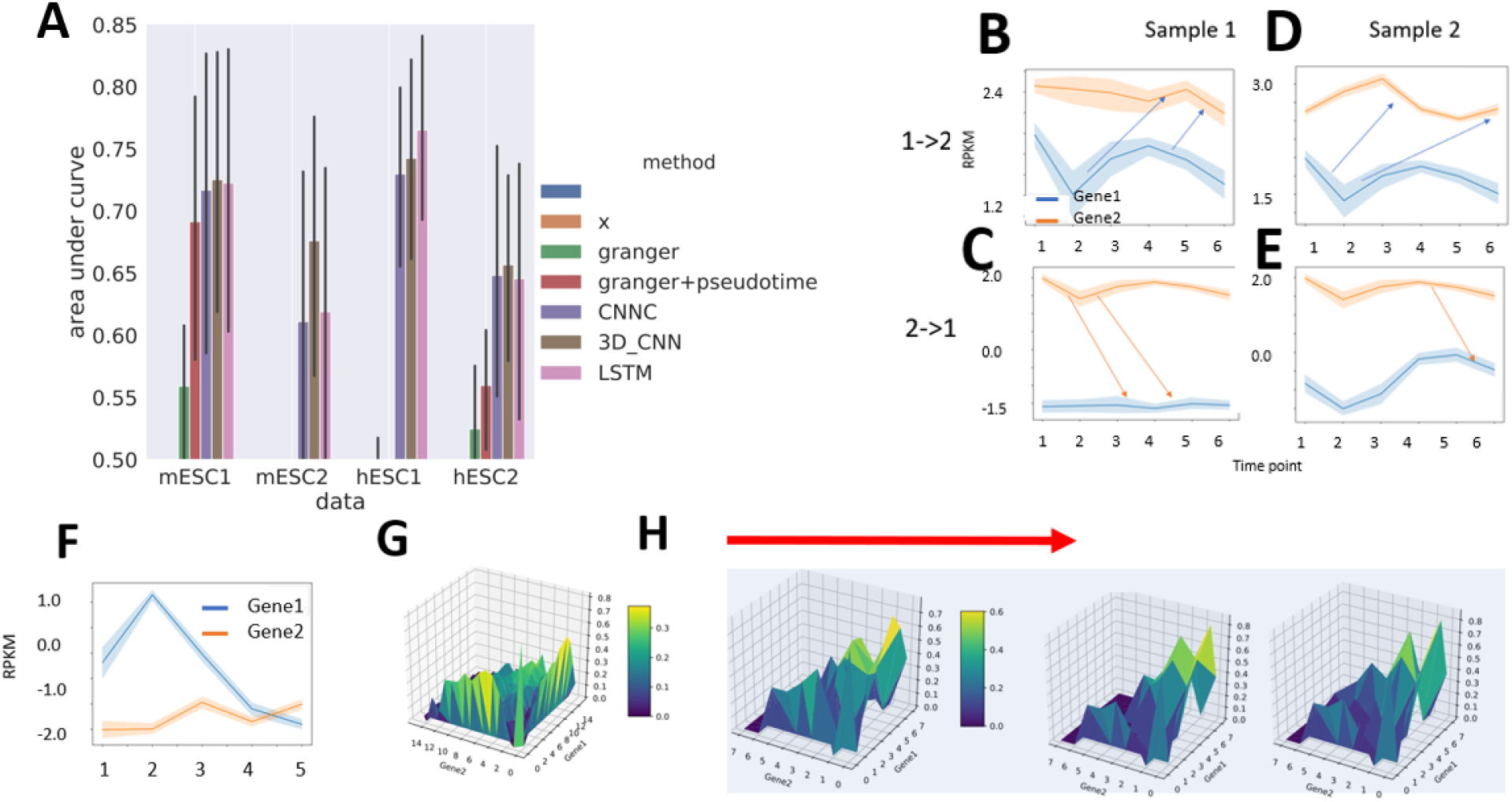
Causality prediction. (A) AUROC of granger causality (with and without pseudo time ordering), CNNC, 3D CNN, and conv-LSTM on TF-target causality prediction tasks in mESC1, mESC2, hESC1, and hESC2 datasets respectively. (B-E) Average gene pair expression over time of 4 pairs that were correctly predicted as gene1->gene2 (top) and 2->1 (bottom) by TDL models in hESC2 dataset. (F-H) The Average gene expression over time (F), and the actual 2D (G), and 3D NEPDFs along with downsampled time point used by TDL (H) for a pair that was correctly predicted as positive by TDL while wrongly predicted as negative by CNNC in hESC1 dataset.

### 3.2 Using TDL to predict TF target genes

Next we tested the ability of TDL models to predict general gene interactions using time series scRNA-Seq data. Here we used the same datasets as in the previous section but changed the task performed by the models so that they attempt to find interactions and not just causal relationships. Specifically, the set of pairs we use for training contains known interacting pairs (positive) which are a TF and its known target and negative (random) pairs which contain a TF and a random gene that is determined to be not a target of that TF. We compared TDL models to several prior methods developed for learning interactions from time series expression data. For all comparisons, we used the labeled training dataset to learn parameters for all supervised models. Specifically, we compared the TDL approaches (3D CNN and Conv-LSTM) with a 2D CNNC (Yuan and Bar-Joseph, 2019) (which is also supervised), two popular unsupervised methods, Pearson correlation (PC) and mutual information (MI) (Song, et al., 2012), and to regression methods for time-course gene expression data. For the regression comparison we used dyngenie3 (Huynh-Thu and Geurts, 2018) which performs random forest regression and uses it to select the key features (other genes) for each gene in two ways, as discussed before (either using the average expression for each time point or applying the method to pseudo-time ordered cells)

Results are presented in **Fig. 3A** (See **Fig. S9** for results of all models). As can be seen, the supervised methods outperformed the unsupervised methods (PC and MI) in all tasks. They were also much better than the regression models and were still lower than the TDL results. In addition, the TDL models outperformed 2D CNNC in three of the four datasets. To gain insight about the information the TDL methods utilize to accurately predict interactions we plotted input samples that were correctly predicted as interacting (label of 1) or not-interacting (0) by TDL. As can be seen in **Fig. 2B and D**, unlike 2D input based methods, TDL methods can take advantage of phase difference between the two genes to correctly determine that they are interacting. While prior methods have also utilized delayed or time shifted interactions (Huynh-Thu and Geurts, 2018), these required explicit choices or search for the duration of the shifted curves. In contrast, TDL were able to infer such shifts without any explicit setting and are flexible adapting to the specific experiments and gene pair shifts. Note also that simple correlation may not be enough to infer interactions. For example, in the hESC2 dataset, the two genes presented in **Fig. 2C and 2E** are highly correlated and yet the TDL methods correctly inferred that they are not interacting. It is well known that gene expression correlation often results from co-regulation of pairs of genes and is not always the result of direct interactions. The fact that TDL models were able to infer such phenomena from training data is a strong indication for their flexibility and ability to focus on relevant information. We also plotted positive samples that were correctly predicted by TDL model (**Fig. 2F-H**) while wrongly assigned as not interacting by the 2D CNNC. We again observe that the models seem to focus on both phase delay between genes along the time axes (**Fig. 2F**), and dynamics among 2D NEPDF in each time point (**Fig. 2H**).

**Fig. 3.**
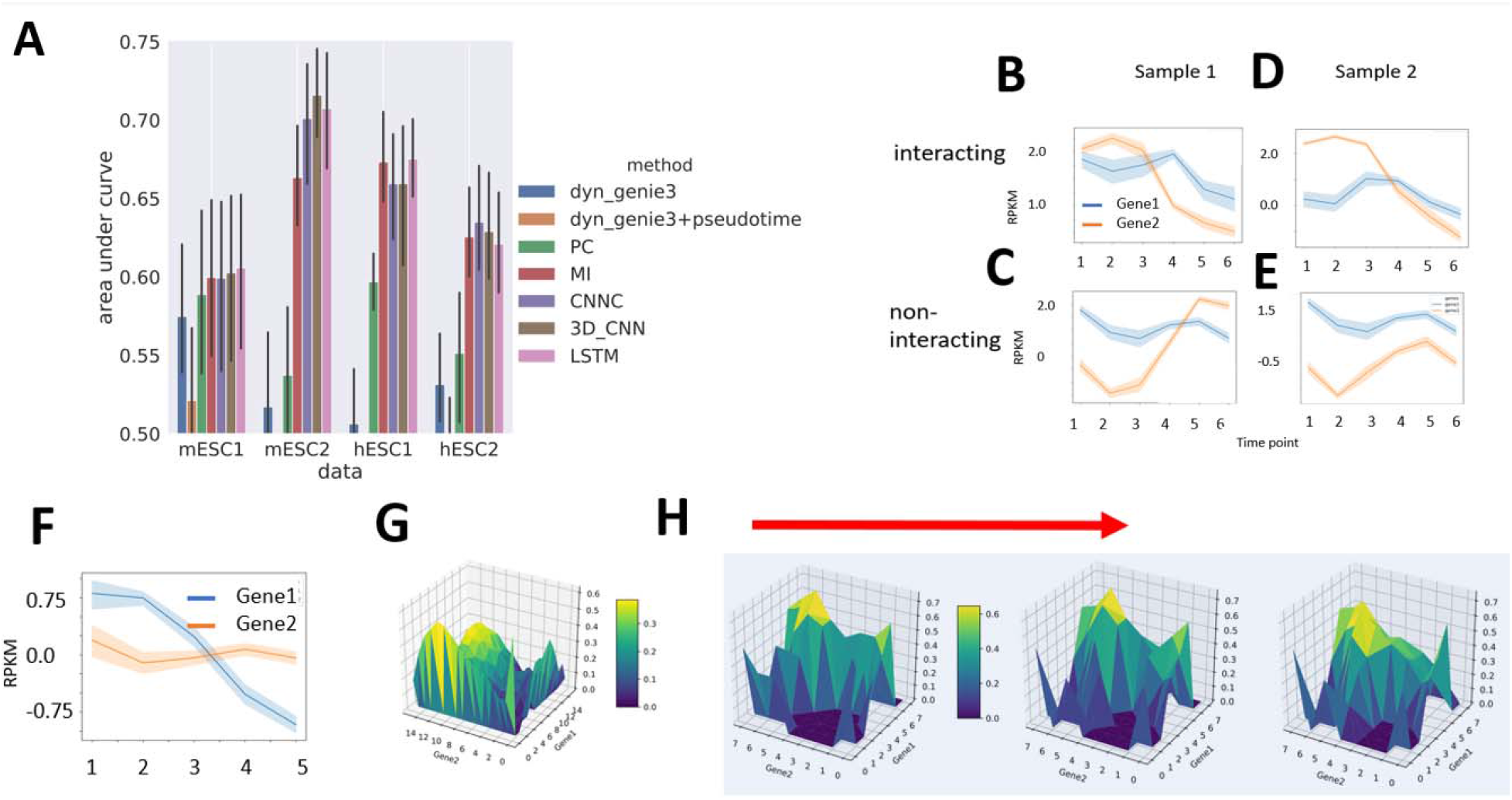
TF target prediction. (A) AUROC of dyngenie3, dyngenie3 with pesudotime ordering by Monocle3, Pearson correlation (PC), mutual information (MI), CNNC, 3D CNN, and conv-LSTM on TF-target prediction in mESC1, mESC2, hESC1, and hESC2 datasets respectively. (B-E) Average gene pair expression along with time point of typical samples that were correctly predicted as interacting gene pairs and non-interacting gene pairs by TDL models in hESC2 dataset. (F-H) Average gene pair expression along with time point (F), overall 2D NEPDF used by CNNC (G), and 3D NEPDF along with downsampled time point used by TDL (H) of a sample that was correctly predicted as positive by TDL while wrongly predicted as negative by CNNC in hESC1 dataset.

To explore additional interactions beyond TF-gene relationships we next selected the top 1,000 highly expressed genes from the MESC2 dataset, and used a model trained using TF-target pairs to score all possible gene pairs among these 1,000 genes. Manual inspection showed that among the top 10 predicted pairs (none of which include TFs used for training), five are supported by recent studies (**Tab. 2**). For example, ‘ndufa4’ is known to be upregulated by ‘apoe4’ (Nuriel, et al., 2017), and ‘igfbp2’ and ‘hist1h2ao’ are both involved in a pathway activated for Alcohol and Stress Responses (Luo, et al., 2018).

**Table 2.**
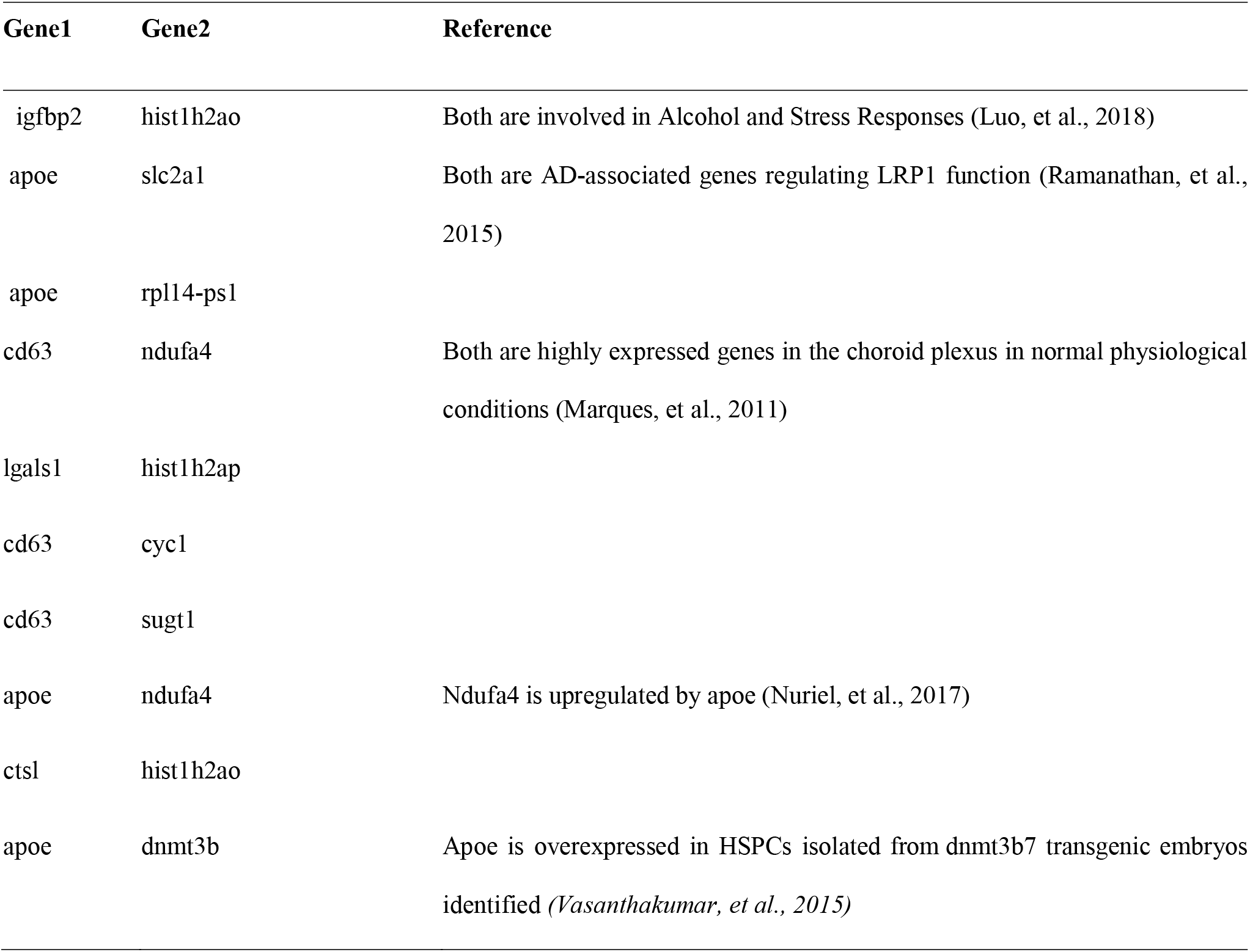
Top predicted gene pairs based on conv-LSTM

| Gene1 | Gene2 | Reference |
| --- | --- | --- |
| igfbp2 | hist1h2ao | Both are involved in Alcohol and Stress Responses (Luo, et al., 2018) |
| apoe | slc2a1 | Both are AD-associated genes regulating LRP1 function (Ramanathan, et al., 2015) |
| apoe | rpl14-ps1 |  |
| cd63 | ndufa4 | Both are highly expressed genes in the choroid plexus in normal physiological conditions (Marques, et al., 2011) |
| lgals1 | hist1h2ap |  |
| cd63 | cyc1 |  |
| cd63 | sugt1 |  |
| apoe | ndufa4 | Ndufa4 is upregulated by apoe (Nuriel, et al., 2017) |
| ctsl | hist1h2ao |  |
| apoe | dnmt3b | Apoe is overexpressed in HSPCs isolated from dnmt3b7 transgenic embryos identified (Vasanthakumar, et al., 2015) |

**Tab. S1** presents the run time for the different NN methods we compared. As can be seen, while TDL models improve on 2D CNNC, their run time is much larger (when using the optimal input resolution for TDL (8×8) and for 2D (16×16)).

### 3.3 Using TDL for assigning functions to genes

TDL can also be used to predict indirect or functional gene relationships. For example, there are hundreds of genes involved in cell cycle related activities. All of them share the same functional annotation though most do not interact. We then used TDL to identify new ‘functions for genes based on known genes for that function. For this we focused on the following categories: cell cycle, rhythm, immune and proliferation.

We downloaded the known gene sets from GSEA (Subramanian, et al., 2005) for cell cycle, rhythm, immune and proliferation. We used 2/3 of the genes for each function for training and the remaining genes for testing. For the function prediction tasks we used a mouse brain time-course single cell data since it contains many more cells (21K cells) than the mESC and hESC datasets.

As can be seen in **Fig. 4A-D**, for cell cycle function assignment, conv-LSTM achieves an AUROC of 0.72, compared to 0.68 for CNNC and 0.67 for 3D CNN. Conv-LSTM also outperforms other methods significantly for all functions. To explore the reason for TDL’s improvement over CNNC, we selected two positive samples that were correctly predicted by conv-LSTM, while 2D CNNC predicts one correctly as cell cycle (**Fig. 4E, F**) and the other, incorrectly, as not (**Fig. 4G, H**). We then plotted the 2D and 3D NEPDF used by the two methods. As can be seen, for samples correctly predicted by both conv-LSTM and CNNC, both 2D and 3D NEPDF display cyclic-like patterns. However, for the ones on which 2D CNNC makes mistake we observe such dynamics for only a subset of the time points (either due to noise or sampling size) which can still be identified by conv-LSTM, but obscures the average computed by 2D CNNC.

**Fig. 4.**
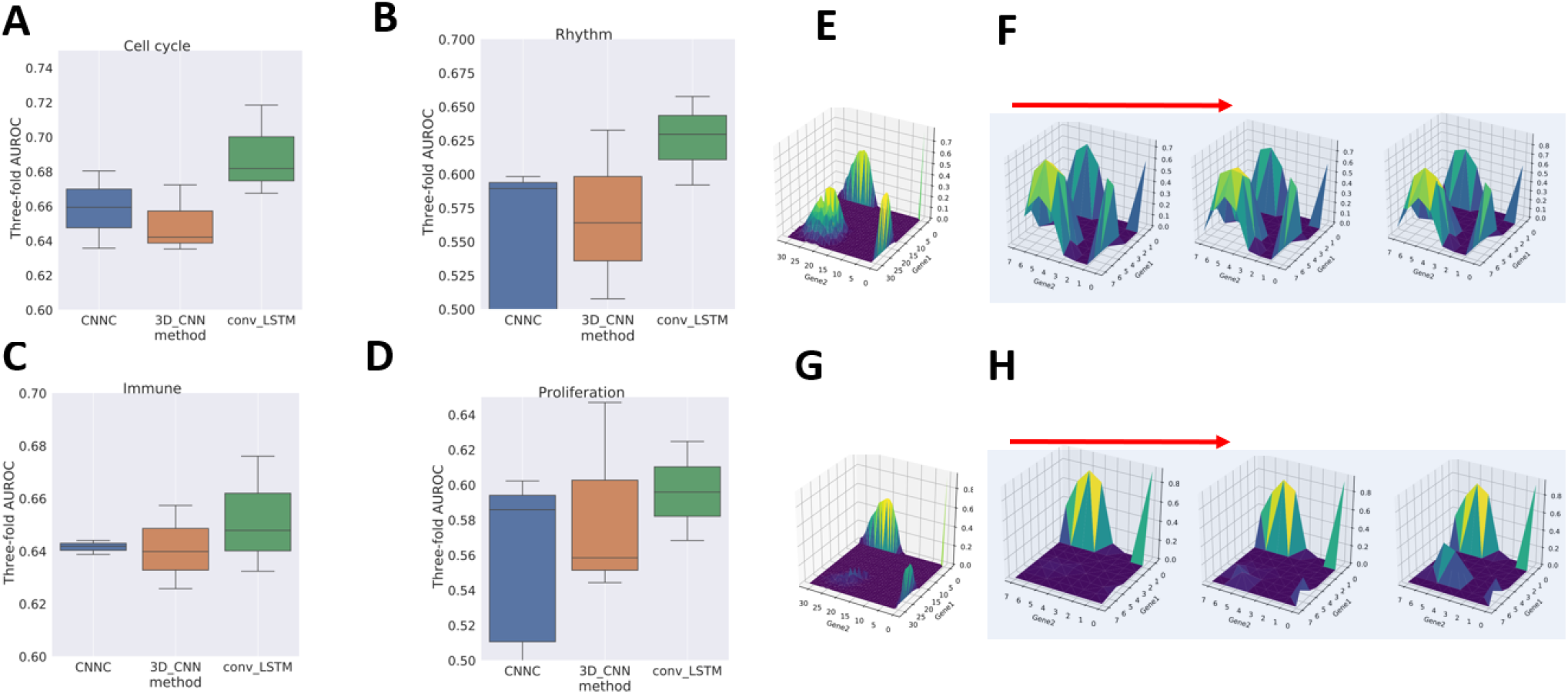
Function assignment. (A-D) AUROC of CNNC, 3D CNN and conv-LSTM on the function prediction task for cell cycle, rhythm, immune and proliferation genes respectively. (E, F) The 2D NEPDF used by CNNC and 3D NEPDF used by conv-LSTM for a positive pair which both CNNC and conv-LSTM correctly classified. (G, H) 2D and 3D NEPDF for a pair that was correctly classified by conv-LSTM while incorrectly classified by CNNC.

## 4 Discussion

Several computational methods have been developed for inferring interactions and causality from time series gene expression data. Most of these methods were developed for bulk data and while some have been also applied to time series scRNA-Seq data, such application is not straightforward. First, in scRNA-Seq analysis there are many more expression profiles when compared to bulk datasets. Second, the ordering of the single cells is much less clear when compared to time series bulk data. Finally, various noise and drop out issues make scRNA-Seq analysis more challenging.

To address these problems we developed Time-course Deep Learning (TDL) methods for the analysis of time series scRNA-Seq data. TDL models rely on encoding for gene expression data as images or a series of images. Each image captures the histogram of a pair of genes for a specific time point and the series of images capture how such histograms evolve over time. Training TDL with known positive and negative pairs lead to models that can infer interactions from such inputs and can be applied to all unknown pairs to predict interactions, causality and function.

We tested TDL using 5 time series scRNA-Seq datasets ranging in size from 700 to 21,000 cells. TDL models were used to perform several tasks including predicting the direction of an interaction, predicting gene-gene interaction (targets of TFs), and predicting the function of a gene. In most of cases TDL models outperform methods developed for bulk data analysis and methods developed for the analysis of static scRNA-Seq data. In addition, as we showed, TDL can accurately predict novel interactions. Thus, TDL can be used to greatly reduce experimental time, costs and effort when studying interactions in specific cell types. Using TDL we can profile a small subset of TFs in these cells (using bulk ChIP-Seq experiments) and combine the outcome with time series scRNA-Seq to accurately predict targets for TFs that were not experimentally profiled.

While TDL worked well for the datasets we looked at, there are still several ways in which they can be improved. Determining the optimal dimension for the input NEPDF, which should be a function of the number of cells and the number of time point, is a challenge. In addition, the current architecture is dependent on the number of time points and so the same model cannot be applied to a different dataset, even for the same condition, if the number of time points do not match.

Our TDL models are implemented in Python. Complete code and sample data is available from the supporting website.

## References

Bar-Joseph, Z., Gitter, A. and Simon, I. Studying and modelling dynamic biological processes using time-series gene expression data. Nat Rev Genet 2012;13(8):552–564.

Cao, J., et al. The single-cell transcriptional landscape of mammalian organogenesis. Nature 2019;566(7745):496–502.

Chu, L.F., et al. Single-cell RNA-seq reveals novel regulators of human embryonic stem cell differentiation to definitive endoderm. Genome Biol 2016;17(1):173.

Finkle, J.D., Wu, J.J. and Bagheri, N. Windowed Granger causal inference strategy improves discovery of gene regulatory networks. Proc Natl Acad Sci U S A 2018;115(9):2252–2257.

Hochreiter, S. The vanishing gradient problem during learning recurrent neural nets and problem solutions. International Journal of Uncertainty, Fuzziness and Knowledge-Based Systems 1998;6(02):107–116.

Hochreiter, S. and Schmidhuber, J. Long short-term memory. Neural computation 1997;9(8):1735–1780.

Huynh-Thu, V.A. and Geurts, P. dynGENIE3: dynamical GENIE3 for the inference of gene networks from time series expression data. Sci Rep 2018;8(1):3384.

Huynh-Thu, V.A., et al. Inferring regulatory networks from expression data using tree-based methods. PLoS One 2010;5(9).

Kim, S., et al. A Granger causality measure for point process models of ensemble neural spiking activity. PLoS Comput Biol 2011;7(3):e1001110.

Klein, A.M., et al. Droplet barcoding for single-cell transcriptomics applied to embryonic stem cells. Cell 2015;161(5):1187–1201.

Luo, J., et al. Integrating Genetic and Gene Co-expression Analysis Identifies Gene Networks Involved in Alcohol and Stress Responses. Front Mol Neurosci 2018;11:102.

Marbach, D., et al. Wisdom of crowds for robust gene network inference. Nat Methods 2012;9(8):796–804.

Marques, F., et al. Transcriptome signature of the adult mouse choroid plexus. Fluids Barriers CNS 2011;8(1):10.

Mayer, C., et al. Developmental diversification of cortical inhibitory interneurons. Nature 2018;555(7697):457–462.

Mikolov, T., et al. Recurrent neural network based language model. In, Eleventh annual conference of the international speech communication association. 2010.

Min, S., Lee, B. and Yoon, S. Deep learning in bioinformatics. Brief Bioinform 2017;18(5):851–869.

Nuriel, T., et al. The Endosomal-Lysosomal Pathway Is Dysregulated by APOE4 Expression in Vivo. Front Neurosci 2017;11:702.

Petropoulos, S., et al. Single-Cell RNA-Seq Reveals Lineage and X Chromosome Dynamics in Human Preimplantation Embryos. Cell 2016;165(4):1012–1026.

Qian, J., et al. Beyond synexpression relationships: local clustering of time-shifted and inverted gene expression profiles identifies new, biologically relevant interactions. J Mol Biol 2001;314(5):1053–1066.

Ramanathan, A., et al. Impaired vascular-mediated clearance of brain amyloid beta in Alzheimer’s disease: the role, regulation and restoration of LRP1. Front Aging Neurosci 2015;7:136.

Schulz, M.H., et al. DREM 2.0: Improved reconstruction of dynamic regulatory networks from time-series expression data. BMC Syst Biol 2012;6:104.

Schulz, M.H., et al. Reconstructing dynamic microRNA-regulated interaction networks. Proc Natl Acad Sci U S A 2013;110(39):15686–15691.

Semrau, S., et al. Dynamics of lineage commitment revealed by single-cell transcriptomics of differentiating embryonic stem cells. Nat Commun 2017;8(1):1096.

Shin, D., et al. Multiplexed single-cell RNA-seq via transient barcoding for simultaneous expression profiling of various drug perturbations. Sci Adv 2019;5(5):eaav2249.

Soldatov, R., et al. Spatiotemporal structure of cell fate decisions in murine neural crest. Science 2019;364(6444).

Song, L., Langfelder, P. and Horvath, S. Comparison of co-expression measures: mutual information, correlation, and model based indices. BMC Bioinformatics 2012;13:328.

Specht, A.T. and Li, J. LEAP: constructing gene co-expression networks for single-cell RNA-sequencing data using pseudotime ordering. Bioinformatics 2017;33(5):764–766.

Stuart, J.M., et al. A gene-coexpression network for global discovery of conserved genetic modules. Science 2003;302(5643):249–255.

Subramanian, A., et al. Gene set enrichment analysis: a knowledge-based approach for interpreting genome-wide expression profiles. Proc Natl Acad Sci U S A 2005;102(43):15545–15550.

Trapnell, C., et al. The dynamics and regulators of cell fate decisions are revealed by pseudotemporal ordering of single cells. Nat Biotechnol 2014;32(4):381–386.

Vasanthakumar, A., et al. Epigenetic Control of Apolipoprotein E Expression Mediates Gender-Specific Hematopoietic Regulation. Stem Cells 2015;33(12):3643–3654.

Wang, Z., et al. DTWscore: differential expression and cell clustering analysis for time-series single-cell RNA-seq data. BMC Bioinformatics 2017;18(1):270.

Xingjian, S., et al. Convolutional LSTM network: A machine learning approach for precipitation nowcasting. In, Advances in neural information processing systems. 2015. p. 802–810.

Yevshin, I., et al. GTRD: a database of transcription factor binding sites identified by ChIP-seq experiments. Nucleic Acids Res 2017;45(D1):D61–D67.

Yuan, Y. and Bar-Joseph, Z. Deep learning for inferring gene relationships from single-cell expression data. Proc Natl Acad Sci U S A 2019.

Zhang, Y., et al. Model-based analysis of ChIP-Seq (MACS). Genome Biol 2008;9(9):R137.

Zou, M. and Conzen, S.D. A new dynamic Bayesian network (DBN) approach for identifying gene regulatory networks from time course microarray data. Bioinformatics 2005;21(1):71–79.

